# Forest wildflowers bloom earlier as Europe warms – but not everywhere equally

**DOI:** 10.1101/2021.09.03.458850

**Authors:** Franziska M. Willems, J. F. Scheepens, Oliver Bossdorf

## Abstract

Some of the most striking biological responses to climate change are the observed shifts in the timing of life-history events of many organisms. Plants, in particular, often flower earlier in response to climate warming, and herbarium specimens are excellent witnesses of such long-term changes. However, in large-scale analyses the magnitude of phenological shifts may vary geographically, and the data are often clustered, and it is thus necessary to account for spatial correlation to avoid geographical biases and pseudoreplication. Here, we analysed herbarium specimens of 20 spring-flowering forest understory herbs to estimate how their flowering phenology shifted across Europe during the last century. Our analyses show that on average these forest wildflowers now bloom over six days earlier than at the beginning of the last century. These changes were strongly associated with warmer spring temperatures. Plants flowered on average of 3.6 days earlier per 1°C warming. However, in some parts of Europe plants flowered earlier or later than expected. This means, there was significant residual spatial variation in flowering time across Europe, even after accounting for the effects of temperature, precipitation, elevation and year. Including this spatial autocorrelation into our statistical models significantly improved model fit and reduced bias in coefficient estimates. Our study indicates that forest wildflowers in Europe strongly advanced their phenology in response to climate change during the last century, with potential severe consequences for their associated ecological communities. It also demonstrates the power of combining herbarium data with spatial modelling when testing for long-term phenology trends across large spatial scales.

## Introduction

Since the industrial revolution anthropogenic global change threatens species and ecosystems. Climate warming in particular can cause shifts in the timing of annual life-history events of plants and animals (Root et al. 2003, Menzel et al. 2006, Cleland et al. 2007). Such phenological changes, including earlier leaf-out or flowering of plants, are some of the most striking large-scale biological responses to ongoing climate change (Cleland et al. 2007). To understand why and how phenology shifts, it is critical to infer which attributes of the environment are the triggers (cues) or proximate causes (drivers) of life cycle events. As their phenology links plants to their environments, changes in the phenology can affect the local persistence and biotic interactions of plants (Inouye 2008, Willis et al. 2008, Wheeler et al. 2015, Cerdeira Morellato et al. 2016). For instance, Willis et al. (2008) found that plant species whose flowering time poorly tracked temperature variation declined in abundance during the last century. Unequal shifts of interacting organisms in trophic interactions can result in phenological “mismatches”, e.g. when the timing of the activity of consumers aligns less well with the availability of their resources, or when the phenology of plants and pollinators shift differently (Renner and Zohner 2018, Visser and Gienapp 2019). Such mismatches can have severe demographic and evolutionary consequences (reviewed e.g. in (Renner and Zohner 2018, Visser and Gienapp 2019).

When studying phenology changes over time, we should keep in mind that phenology, and magnitudes of phenological responses to climate change, not only vary among species but they also vary in space. At smaller scales, phenology can vary because of microclimatic differences (Hwang et al. 2011; Ward et al. 2018; Willems et al. 2021), and at larger scales both (baseline) phenology as well as phenological responses are expected to vary because of macroclimatic variation, because the magnitudes of climatic changes differ geographically (Klein Tank et al. 2002, IPPC 2019), and because phenological sensitivities to cues such as temperature may differ between regions (Riihimäki and Savolainen 2004, Zohner and Renner 2014, Prevéy et al. 2017, Zohner et al. 2020, Kopp et al. 2020). Robust studies on phenology and climate change therefore require a larger-scale perspective, with spatial variation and autocorrelation explicitly taken into account. However, many previous studies on plant phenological responses to climate change had a limited geographical scope (Pau et al. 2011).

In this context, herbaria offer unique opportunities because they allow tracking phenology at large temporal as well as spatial scales. Herbarium specimens are usually collected when plants flower, and most herbarium sheets provide collection dates and locations (Fig. 1). With many herbaria dating back to some 200 years, and hundred millions of specimens worldwide, herbaria are a tremendous treasure for studying phenology changes both long-term and large-scale. Previous studies have indeed found strong patterns of long-term phenology changes in herbarium data (Primack et al. 2004, Miller-Rushing et al. 2006, Davis et al. 2015, Willis et al. 2017, Lang et al. 2019, Park et al. 2019, reviewed by Jones and Daehler 2018), and they have also demonstrated that phenology trends estimated from herbarium data are comparable to those from field observations (Davis et al. 2015, Jones and Daehler 2018, Miller et al. 2021). However, almost all previous studies were done in the US, and there has been little work so far on herbaria and plant phenology in Europe (but see Robbirt et al. 2011, Molnar et al. 2012 and Diskin et al. 2012). Most previous studies also did not consider geographic variation in phenology and spatial correlation of herbarium samples (but see Matthews and Mazer 2016, Park et al. 2019, Kopp et al. 2020).

**Figure 1.**
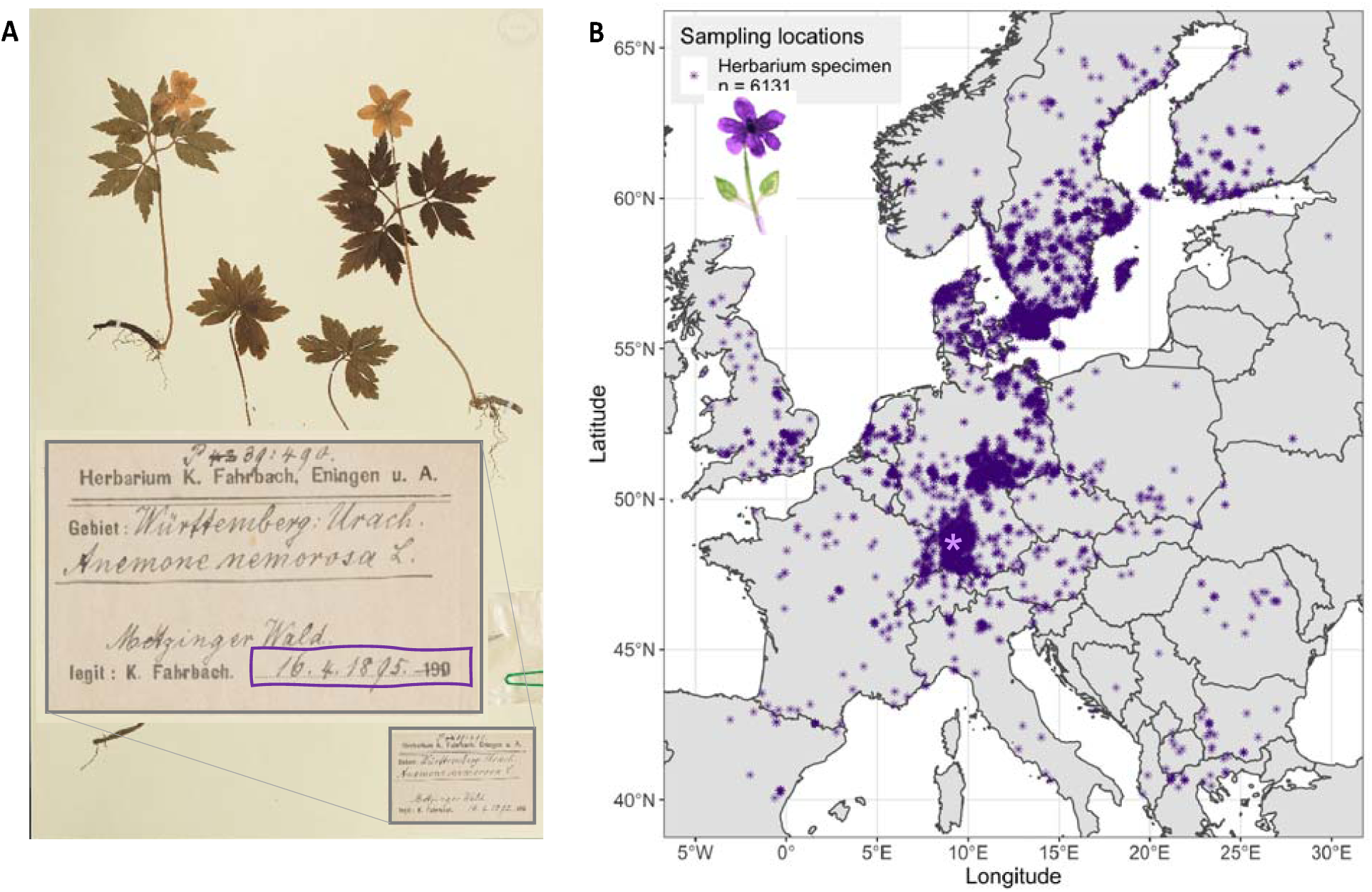
(A) Example of an herbarium specimen, with the collection date and location on the herbarium label. This *Anemone nemorosa* was flowering on April 16 (DOY = 107) in 1895, and it was collected in the “Metzinger Wald” forest close to Tübingen (lighter purple point in the map). (B) Sampling locations of the 6131 herbarium specimens included in our analyses.

In Europe, climatic conditions vary substantially across the ranges of many plant species, especially from north to south, and not only the overall timing of phenological events but also phenological responses (i.e. sensitivities to cues or drivers) may differ across this latitudinal gradient. For a similar climatic gradient in the eastern US, Park et al. (2019) found that long-term phenological responses estimated from herbarium specimens substantially differed among climatic zones, with greater mean climate sensitivities, as well as greater among-species variability in sensitivities, in the warm and mixed-temperate climatic regions than in the cool-temperate northeast and the Appalachians. Similarly, for the Pacific North West region of North America Kopp et al. (2020) found that sensitivity to temperature was greater at low elevations and in the maritime (western) regions. Another problem with large-scale herbarium data is that they are often, for historical reasons, strongly clustered, i.e. specimens are more frequently collected where collectors live, and around academic institutions. However, when modelling across a spatial range, standard methods such as linear regression ignore the spatial dependency between sampling locations and treats all data points as independent. This assumption is very likely not correct, since the proximity of spatial locations is usually related to their environmental similarity (Tobler 1970), and as explained above, this is certainly true for climatic conditions. Ignoring spatial dependency thus results in pseudoreplication, and it can strongly bias model results. The solution to this, spatial modeling with explicit incorporation of spatial structure and thus spatial autocorrelation, is computationally challenging, and it has therefore hardly been used in analyses of herbarium data. However, recent advances in statistical methods now allow to model such spatial data, e.g. using stochastic partial differential equations (SPDE) and integrated nested Laplac approximations (INLA) as implemented in the R package *R-INLA* (Rue et al. 2017, Bakka et al. 2018), and it is therefore possible to take the next step in herbarium studies and analyse large-scale phenology in relation to climate change in a spatially explicit framework.

Here, we analysed long-term and large-scale trends in the flowering time of 20 common forest understory wildflowers, and their relationships with climate change, across Europe, using over a century of herbarium data. We focused on early-flowering understory plants, because they have a very distinct phenology, with a critical time window for flowering before the leaf-out of deciduous trees. Because of this, they may be particularly sensitive to climate change and phenology shifts. Forest understory plants may also be exposed differently to climate change because climate warming is buffered under forest canopies (De Frenne et al. 2019). In our analyses, we employed R-INLA (Rue et al. 2009, 2017, Bakka et al. 2018) to account for spatial clustering and autocorrelation of climate and phenology data. We asked two main questions: (A) Did forest understory plants advance their flowering phenology during the last ∼100 years? (B) If yes, are these phenological shifts associated with climate change in Europe? We answered both questions with or without accounting for spatial correlation in the statistical models, and thus also addressed the question of how important doing this was for the results and conclusions of our study.

## Methods

### Phenological data

We mined three large German herbaria and the Global Biodiversity Information Facility (GBIF) for all European specimens of 20 common spring-flowering forest understory herbs (see Table S1). The three herbaria were at the University of Tübingen (international herbarium code TUB), University of Jena (JE) and at the State Museum of Natural History in Stuttgart (STU). Our criteria for including herbarium specimens were that: (i) they had flowers, and that open flowers represented at least 50% of the reproductive structures, (ii) they had an exact collection date and (iii) information on the sampling location that we could use to estimate GPS coordinates, and (iv) they were collected in Europe. In addition, we obtained all digital specimens of the same 20 species from GBIF (GBIF 2020) that were from Europe and also had (i) an exact collection date and (ii) GPS coordinates of the sampling location, using the *rgbif* package (Chamberlain and Boettiger 2017) in R (R Core Team 2008). This resulted in an initial 3930 specimens from the three herbaria and 3511 specimens from GBIF, with the collection years ranging from 1807 – 2017. However, since reliable, gridded climate data were not available before 1901 we decided to restrict our analyses to data from 1901 onwards. Moreover, because there were only very few specimens from outside of these limits, we truncated our data to 40 to 65 degrees northern latitude and −5 to 30 degrees longitude, covering a broad geographic area in mainly Central and Northern Europe, but also Western and South-Eastern Europe (Fig. 1A). We further discarded all specimens with dates outside of the normal flowering range of our 20 study species (before day of the year (DOY) 50 and after DOY 200), because we suspected these to be recording mistakes. Also, the GBIF data contained unusually many specimens from May 1 and June 1 (DOYs 121 and 152, respectively), which strongly indicated that they were from specimens without exact collection dates that were arbitrarily assigned to the first day of a month, and we excluded these data from our analyses. Lastly, we discarded six datapoints for which the assigned elevation value was below −10 m. Our final set of phenology data contained 6131 herbarium specimens, with 46 to 600 records per species (Table S1).

### Climate and elevation data

For associating plant phenology variation with long-term temporal and spatial variation in climate, we used gridded estimates of historic monthly air temperature (°C) and precipitation (mm) that were available for 1901–2017 and with a 0.5° × 0.5° grid resolution from the Climate Research Unit (CRU, https://crudata.uea.ac.uk; (Harris et al. 2020), version *cru_ts4.04*). We used these data to calculate mean winter (December – February) and spring (March – May) temperatures, as well as annual precipitation values for each year and grid cell. Each herbarium specimen was then assigned to a specific set of values of these three climate variables, based on its collection year and the geographic grid cell it was located in, using custom-made scripts in *python* (Van Rossum and Jr. Drake 2009). We also estimated the elevation a.s.l. of each herbarium specimen using the *raster* package in R (Hijmans 2020).

### Statistical analyses

Our statistical analyses generally had a two-step logic, relating to the two main questions of our study. We first tested for overall phenological shifts, i.e. temporal trends in flowering time, across our 20 study species, using a simpler statistical model (model A), and we then tested for phenology-climate associations with a more complex model B (details below). Both models were run with and without accounting for spatial correlation.

To test for temporal trends in flowering time (model A) we modelled flowering phenology during the last 120 years as a function of the year of collection, while accounting for the effects of elevation and species. Model A was specified as:

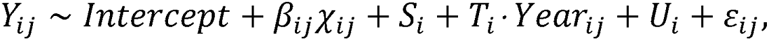

where *Y_ij_* is the day of flowering of herbarium specimen *i* of species *j*, χ*_ij_* is a vector containing all covariates (model A: collection year and elevation) as linear fixed effects, *β_ij_* is the vector of estimated parameters (regression slopes), 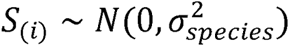 is the species random intercept, 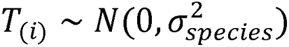 the species random slopes, both with a Gaussian distribution, and ε*_ij_* ∼ *N*(0, *σ*^2^) the residuals. The species random intercept allows species to differ in their mean flowering times, and the species random slope means that temporal trends can be species-specific, e.g. because species respond differently to climate change. U_(*ij*)_ ∼ *N* (0, *Ω*) represents the spatial structure (see below) that is additionally included as a random effect in the models accounting for spatial correlation. In model A, the slope of the linear relationship between the collection dates (= DOY of flowering) of specimens and their collection year is the formal test for long-term phenological shifts.

To test for phenology-climate associations (model B) we additionally included spring temperature, winter temperature and precipitation, plus the interactions between spring temperature (which is usually considered a key driver of spring phenology) and all other variables into the model described above (see Table 1 for more detailed explanations of the variables, and their expected effects on phenology). χ*_ij_* again includes all these covariates and *β_ij_* are their respective effects, i.e. regression slopes We thus modified the model equation to:

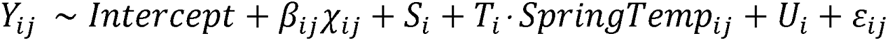

In model B the slopes of the linear relationships between the collection dates (= DOY of flowering) of specimens and the temperature or precipitation at the corresponding location and year estimates the sensitivities of phenology to climate changes. Here, the species random slopes are the species-specific shifts with temperature (*T*_(*i*)_) accounting for the fact that some species might be more temperature sensitive than others. As for model A, we also fitted model B with and without including the spatial structure *U_ij_*.

**Table 1.**
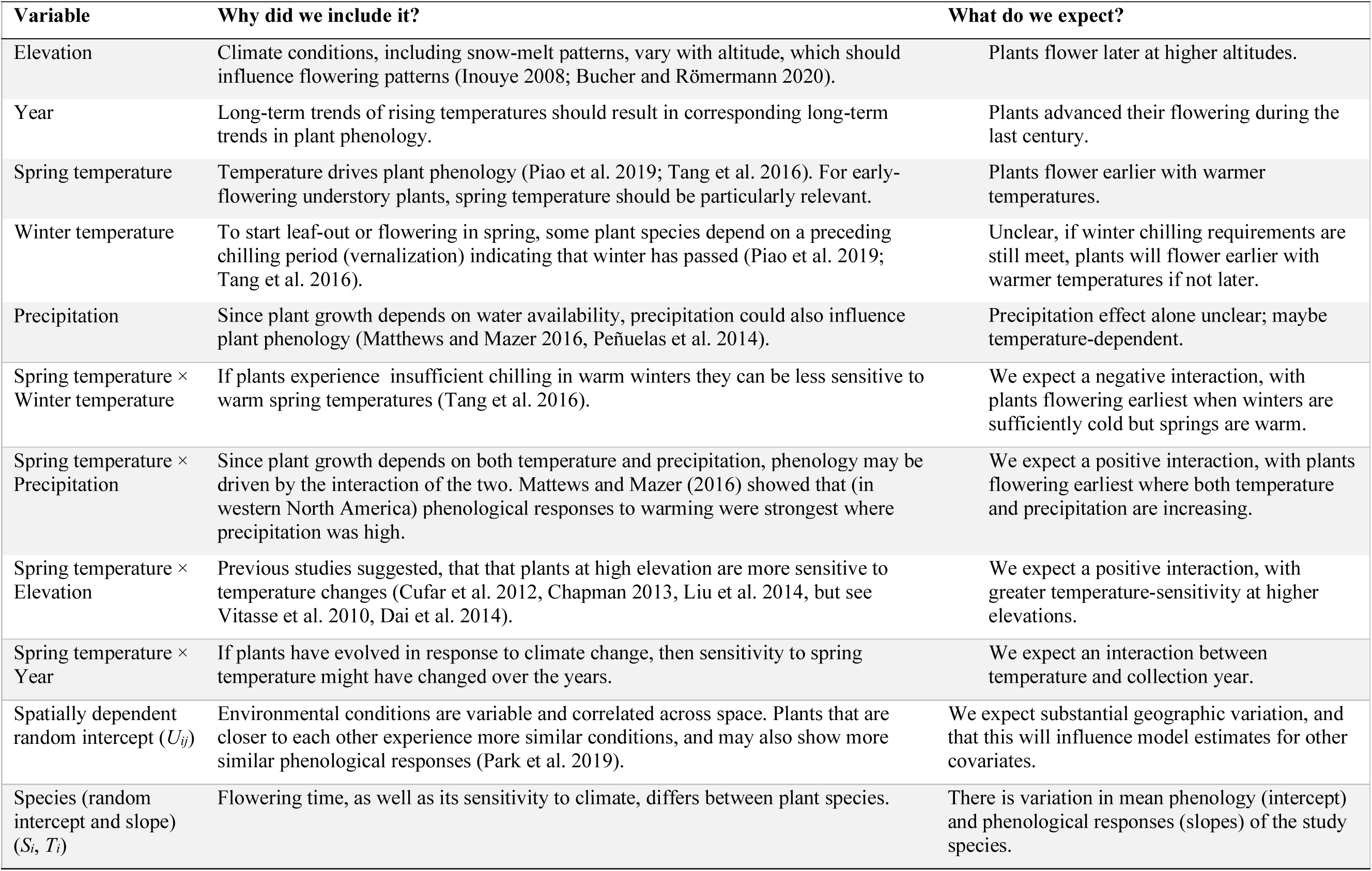
All explanatory variables (fixed and random effects) that were included in our analyses, together with the reasonings for including them, and their expected effects on plant phenology. Model A included elevation, year and the two random factors, model B also the climate variables, and the interactions of spring temperature with the other covariates.

| Variable | Why did we include it? | What do we expect? |
| --- | --- | --- |
| Elevation | Climate conditions, including snow-melt patterns, vary with altitude, which should influence flowering patterns (Inouye 2008; Bucher and Römermann 2020). | Plants flower later at higher altitudes. |
| Year | Long-term trends of rising temperatures should result in corresponding long-term trends in plant phenology. | Plants advanced their flowering during the last century. |
| Spring temperature | Temperature drives plant phenology (Piao et al. 2019; Tang et al. 2016). For early-flowering understory plants, spring temperature should be particularly relevant. | Plants flower earlier with warmer temperatures. |
| Winter temperature | To start leaf-out or flowering in spring, some plant species depend on a preceding chilling period (vernalization) indicating that winter has passed (Piao et al. 2019; Tang et al. 2016). | Unclear, if winter chilling requirements are still met, plants will flower earlier with warmer temperatures if not later. |
| Precipitation | Since plant growth depends on water availability, precipitation could also influence plant phenology (Matthews and Mazer 2016, Peñuelas et al. 2014). | Precipitation effect alone unclear; maybe temperature-dependent. |
| Spring temperature × Winter temperature | If plants experience insufficient chilling in warm winters they can be less sensitive to warm spring temperatures (Tang et al. 2016). | We expect a negative interaction, with plants flowering earliest when winters are sufficiently cold but springs are warm. |
| Spring temperature × Precipitation | Since plant growth depends on both temperature and precipitation, phenology may be driven by the interaction of the two. Matthews and Mazer (2016) showed that (in western North America) phenological responses to warming were strongest where precipitation was high. | We expect a positive interaction, with plants flowering earliest where both temperature and precipitation are increasing. |
| Spring temperature × Elevation | Previous studies suggested, that that plants at high elevation are more sensitive to temperature changes (Cufar et al. 2012, Chapman 2013, Liu et al. 2014, but see Vitasse et al. 2010, Dai et al. 2014). | We expect a positive interaction, with greater temperature-sensitivity at higher elevations. |
| Spring temperature × Year | If plants have evolved in response to climate change, then sensitivity to spring temperature might have changed over the years. | We expect an interaction between temperature and collection year. |
| Spatially dependent random intercept ($U_{ij}$) | Environmental conditions are variable and correlated across space. Plants that are closer to each other experience more similar conditions, and may also show more similar phenological responses (Park et al. 2019). | We expect substantial geographic variation, and that this will influence model estimates for other covariates. |
| Species (random intercept and slope) ($S_i$, $T_i$) | Flowering time, as well as its sensitivity to climate, differs between plant species. | There is variation in mean phenology (intercept) and phenological responses (slopes) of the study species. |

To estimate spatial dependency, we used integrated nested Laplace approximation (INLA), an approximate Bayesian technique and faster alternative to MCMC methods for fitting Bayesian models (Bakka et al. 2018). A key challenge with spatial models is that the Gaussian random field, the most common tool for capturing spatial dependency, is hard to use with large data. *R-INLA* solves this problem through stochastic partial differential equations (SPDEs) that allow to model Gaussian random fields fast and efficiently, and to handle complex spatial data (Lindgren et al. 2011). The SPDE is the mathematical solution to the Matérn covariance function describing the statistical covariance between values at two different points. The covariance matrix of the Gaussian field is approximated as a Gaussian Markov Random Field (GMRF) using a Matérn covariance structure (Bakka et al. 2018). The GMRF models spatial dependence by defining a neighborhood structure on a mesh that divides the study area (in our case Europe) into non-overlapping triangles (Fig. S1). The data points (in our case sampling locations of herbarium specimens) are then assigned to the adjacent nodes of the mesh according to their proximities (or to only one if they fall directly onto one). This creates an observation matrix for estimating the Gaussian Markov Random Field (Bivand et al. 2015, Cosandey-Godin et al. 2015). The mesh can have different shapes and sizes, and we used the default constrained Delaunay triangulation (a particular way to divide an area into triangles) together with vague priors that have little effect on the posterior distributions of the fixed effects. To select the mesh size, we compared models with different meshes and chose the finest mesh (with a maximum triangle edge length of 20 km and a minimum edge length of 5 km) as it resulted in the lowest DIC/WAIC values. The derived Gaussian Markov Random field is then represented by the term *U_ij_* in the model above, a smooth spatial effect that links observations to spatial locations, with the covariance structure *Ω* estimated via the Matérn correlation. The term *U_ij_* is thus spatially variable and captures spatial patterns not already modelled by the fixed covariates, thereby ensuring that the residuals ε_ij_ are independent. We compared the results of models with and without including *U_ij_*.

To avoid biased parameter estimates because of unequal scales, we fitted covariates in the following forms: year expressed in decades, spring precipitation in mm·10^-1^, elevation in hundred meters [100 m] and spring and winter temperature in degree Celsius [°C]. We also mean-centered all covariates because this estimated the regression slopes of each covariate with all other covariates at their mean values (rather than zero; (Dalal and Zickar 2012), which greatly helped to interpret the results of the regression analysis.

For both models we checked whether the residuals were normally distributed, plotted the distribution of residuals against fitted values and explanatory variables to check for heterogeneity or other patterns in the variances, and we plotted the observed vs fitted data to evaluate model fit and performance (Zuur et al., 2017). All statistical analyses were done in R version 3.6.2 (R Core *INLA* Team 2018) using the *R*□*INLA* package (http://www.r-inla.org see also: Rue et al. 2009, see also: Rue et al. 2009, Lindgren et al. 2011, Bakka et al. 2018).

## Results

### Model validation and spatial correlation

The herbarium data analysed in our study covered a broad geographical range in Europe, but their spatial distribution was heterogenous (Fig. 1), and in addition the flowering time data were spatially correlated up to a distance of around 200 km and 100 km in models A and B, respectively (Fig. S2). If this spatial correlation was not included in the analyses, then the model residuals were clearly non-random in space, especially in model A (Fig. S3), and there were other violations of model assumptions, in particular non-random distribution of residuals in relation to several covariates (Fig. S4 and S5). Including spatial correlation strongly solved these problems. Moreover, models that included spatial correlation also generally had a better fit (see Fig. S6 for a comparison of DIC values and regression parameter estimates of model B with and without spatial correlation), and the fitted values were much closer to the observed values (*r* = 0.78 vs. 0.57 for Model A and *r* = 0.82 vs 0.70 for Model B; Fig. S7). Overall, residual variation was reduced when spatial correlation was accounted for (Fig. S8). Thus, models that explicitly incorporate spatial correlation between data points are not only more statistically sound, but they are also stronger and more informative. In the next sections, we show that taking spatial correlation into account also substantially affects the model estimates answering the main questions of our study.

### Temporal shifts in plant phenology

Overall, the herbarium data indicated that the studied 20 forest understory plants significantly advanced their flowering time during the last century (Table 2, Fig. 2A, B). The estimated advancement of flowering time was −0.56 days per decade (credible interval: −0.74 to −0.39; see Table 2) according to model A with accounting for spatial correlation, and these responses were different from zero (posterior probability > 0.95) for all 20 species. For species-specific residuals see Fig. S9 and for a summary of all hyperparameters see Table S2. The observed phenology shifts corresponded with increasingly warmer spring temperatures during the last century (Fig. 2C). If model A ignored spatial correlation, it severely overestimated the overall magnitude of phenology shifts, with an estimated −1.34 days per decade (CI: −1.69 to −0.98; Table S3, Fig. 2B), i.e. it estimated an average shift of around two weeks during the last century, more than twice as much as in model A with spatial correlation. One reason for this discrepancy was that datapoints from northern vs southern Europe were unevenly distributed in time, with more earlier data from the north, and an overrepresentation of southern data during the last decades (Fig. 2C). When spatial information is ignored in model A, this latitudinal bias thus distorts the estimated shift over time. The opposite is true for the relationship with elevation: in model A with spatial correlation plants flower later at higher altitudes (2.44 days/100m, 95% CI 1.98 to 2.89; Table 2, Fig. 2B), but when spatial correlation is ignored there is no relationship between elevation and flowering time (Table S3, Fig. 2B). Even with including spatial correlation, and after the effects of the covariates year and elevation have been accounted for there is still strong spatial variation in flowering time in model A, with plants from Northern and Eastern Europe flowering up to ∼60 days later than plants from Central and Southern Europe (Fig. 5).

**Table 2.** Model estimates (slopes), with standard deviations and 95% credible intervals, for all variables included in models A and B with spatial autocorrelation.

|  | **Estimate** | **SD** | **95% CI** |
| --- | --- | --- | --- |
| **Model A** |  |  |  |
| Intercept | 136.09 | 4.07 | 128.02, 144.03 |
| Years [Decades] | -0.56 | 0.09 | -0.74, -0.39 |
| Elevation [100 m] | 2.57 | 0.24 | 2.08, 3.02 |
| **Model B** |  |  |  |
| Intercept | 138.62 | 2.52 | 133.57, 143.48 |
| Spring temperature [°C] | -3.61 | 0.22 | -4.04, -3.18 |
| Winter temperature [°C] | -1.05 | 0.13 | -1.31, -0.79 |
| Precipitation [mm/10] | 0.07 | 0.15 | -0.23, 0.37 |
| Elevation [100 m] | 1.42 | 0.21 | 1.00, 1.84 |
| Year [Decade] | -0.22 | 0.09 | -0.40, -0.04 |
| Spring temperature × Year | 0.05 | 0.04 | -0.03, 0.13 |
| Spring temperature × Elevation | 0.06 | 0.05 | -0.04, 0.16 |
| Spring temperature × Precipitation | -0.04 | 0.06 | -0.16, 0.07 |
| Spring temperature × Winter temperature | -0.06 | 0.03 | -0.12, 0.01 |

**Figure 2.**
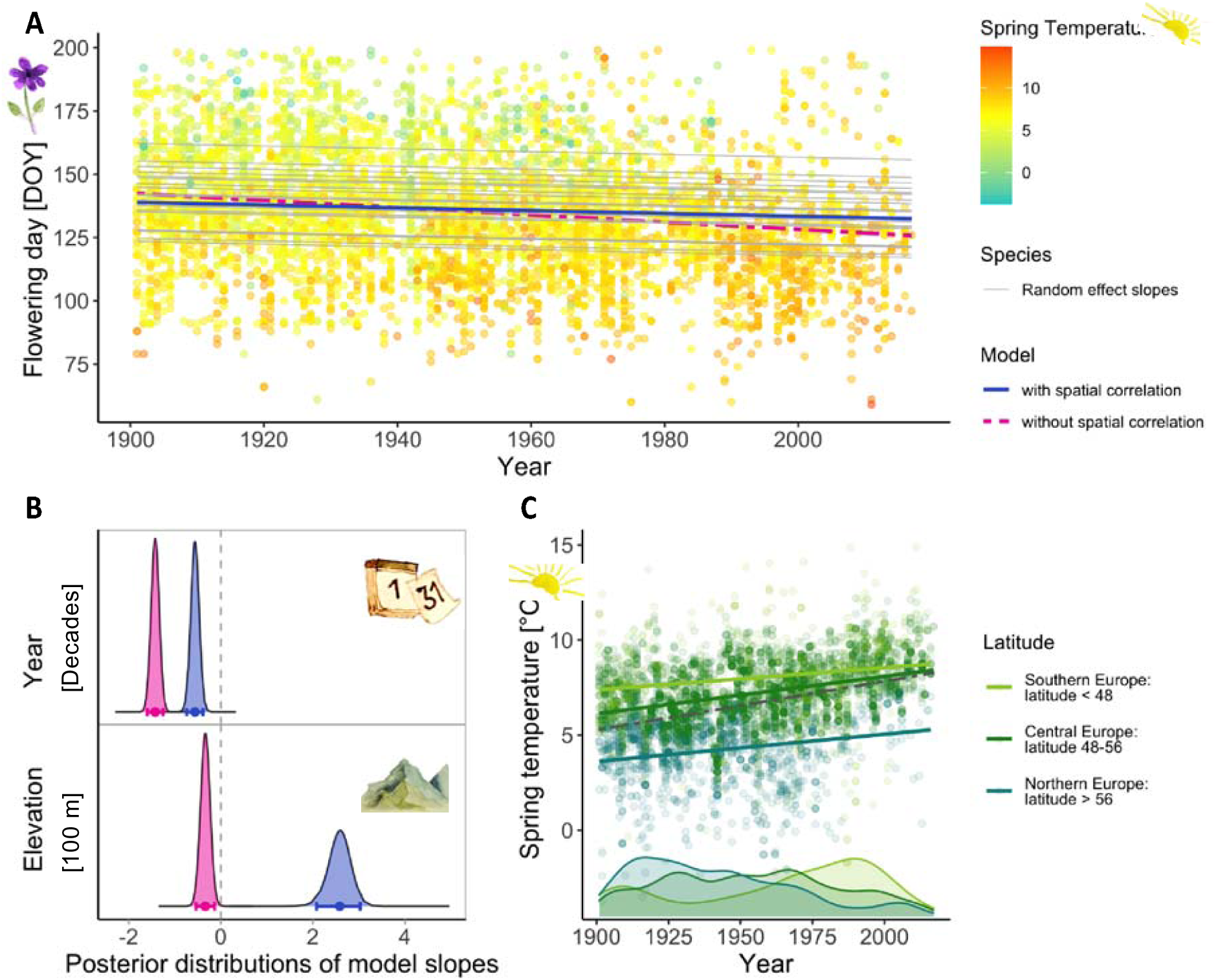
Temporal trends of flowering time and spring temperature over the last century, and the results of model A. (A) Shifts of flowering time since 1901 estimated by model A with spatial correlation (solid blue line) and without spatial correlation (dashed magenta line). With spatial correlation, plants advanced their flowering on average by around six days, and the responses were different from zero (posterior probability > 0.95) for all 20 species (thin grey lines). In the model without spatial correlation the estimated phenology shift is more than twice as large. (B) Differences in parameter estimates (posterior probability distributions) for model for model A without (magenta) and with (blue) spatial correlation. (C) Long-term trends in spring temperature in the locations of the studied herbarium specimens, separately for southern, central and northern European data. The histograms at the bottom show the temporal distributions of these data.

### Relationships with climate change

Across the European sampling locations included in our study, spring temperatures increased during the last century (Fig. 2C), and the phenology of the plants was related to these climatic changes. Overall, plants flowered around 3.6 days earlier per +1°C (Table 2, Fig. 3 and 4). If spatial correlation was not included in model B, the strength of this relationship was overestimated with 5.4 days per +1°C (Fig. 3 and 4). The general temperature-phenology relationship was consistent across the 20 studied species, with negative slopes credibly different from zero (posterior probability > 0.95) for all (Fig. 3).

**Figure 3.**
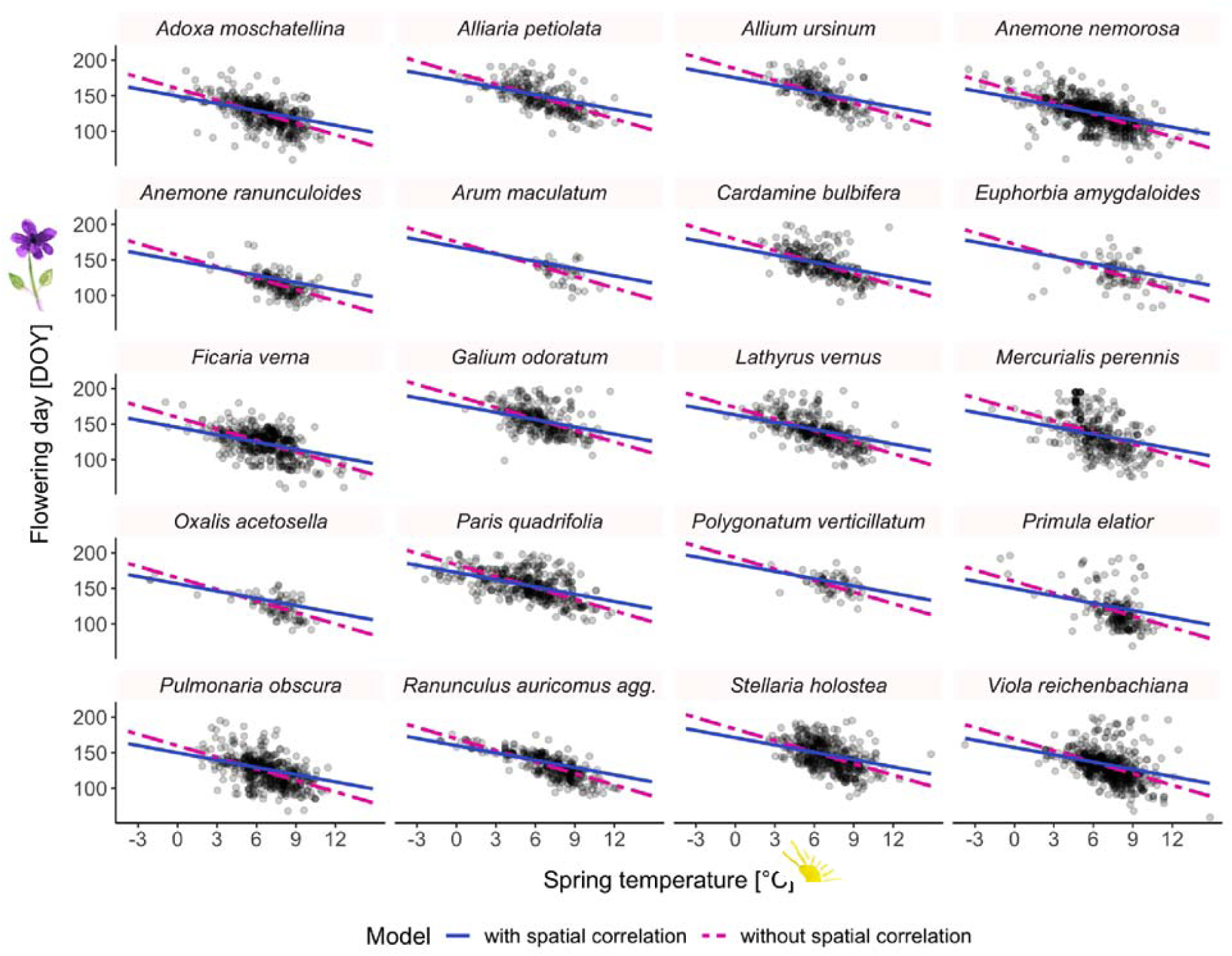
Relationships between the spring (March-May) temperature in the year of collection and the date of collection (= flowering day) of European herbarium specimens of 20 early-flowering forest understory plants. The blue and magenta lines indicate slope estimates from statistical models with and without taking spatial autocorrelation into account, respectively.

Besides the relationship with spring temperature, there was a significant, albeit weaker, relationship with winter temperature, but no relationship with precipitation, in the model B with spatial correlation (Table 2, Fig. 4). There were further relationships of phenology with elevation and the year of sampling (Table 2, Fig. 4). The direction of these results – later flowering at higher altitudes and earlier flowering in more recent specimens – was as in model A, only with smaller effect sizes. This is because both the year of sampling and elevation are systematically related to temperature, so the larger effects in model A are partly temperature effects. None of the interaction terms between spring temperature and the other covariates were significant (Table 2). Ignoring the spatial locations of specimens substantially affected also these parameter estimates: in model B without spatial correlation the relationship with elevation was underestimated, whereas the relationship with winter temperature was lost, and there was now a relationship with precipitation, and several significant interactions between covariates (Table S3, Fig. 4).

**Figure 4.**
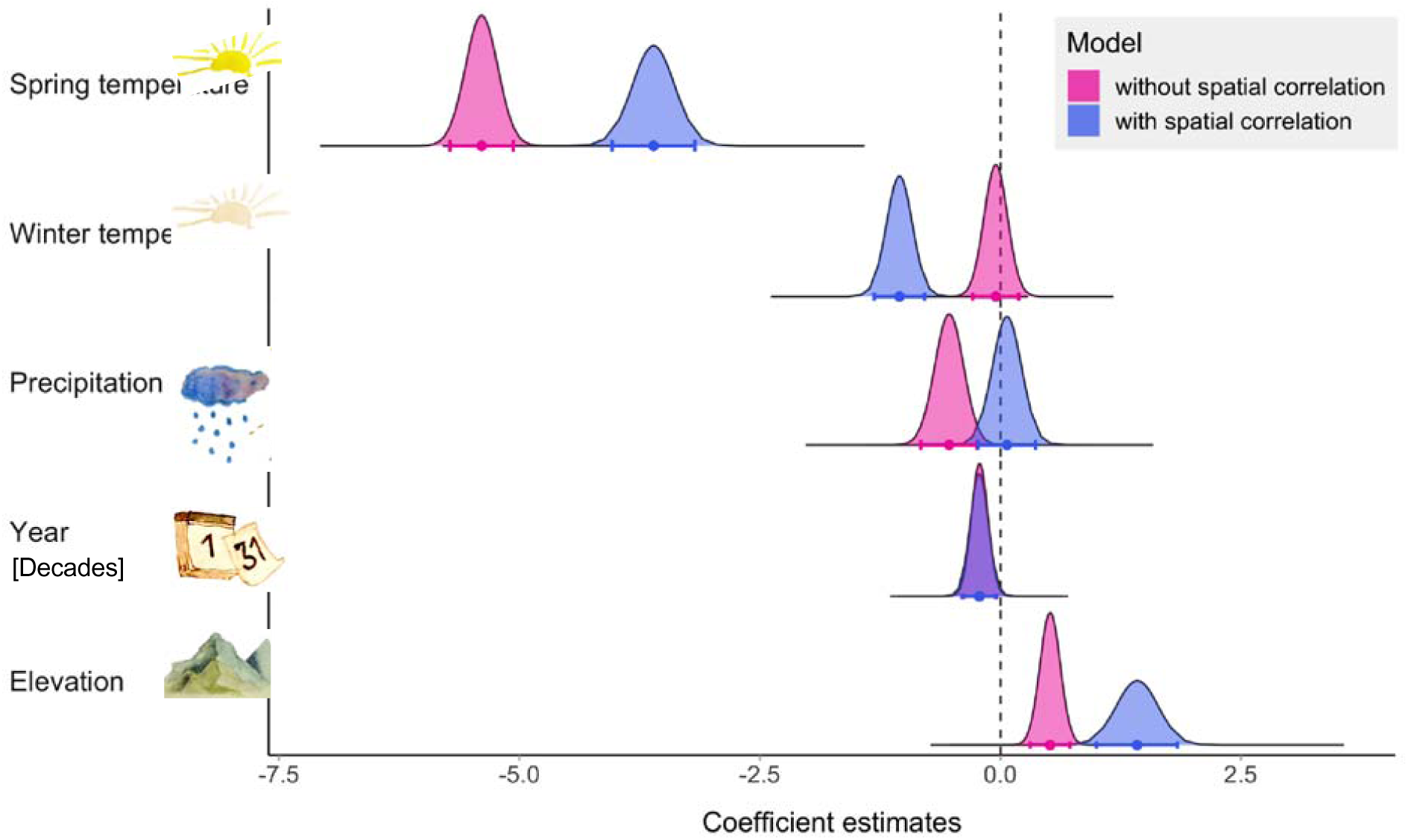
Model coefficient estimates for relationships between different covariates (climate in the year of collection, year of collection, elevation of collection site) and the date of collection (= flowering time) of herbarium specimens of 20 forest wildflowers in Europe. The blue vs. mangenta curves show the differences between the parameter estimates (posterior probability distributions) from model B with and without taking spatial autocorrelation into account.

As in model A, there was significant spatial variation in flowering time after the covariates and their interactions had been accounted for (Fig. 5, right panel). Although the residual spatial correlation was clearly much less and more small-scale than in model A, there were still several regions with clustering of positive or negative residuals, showing the importance of incorporating spatial correlation also in model B.

**Figure 5.**
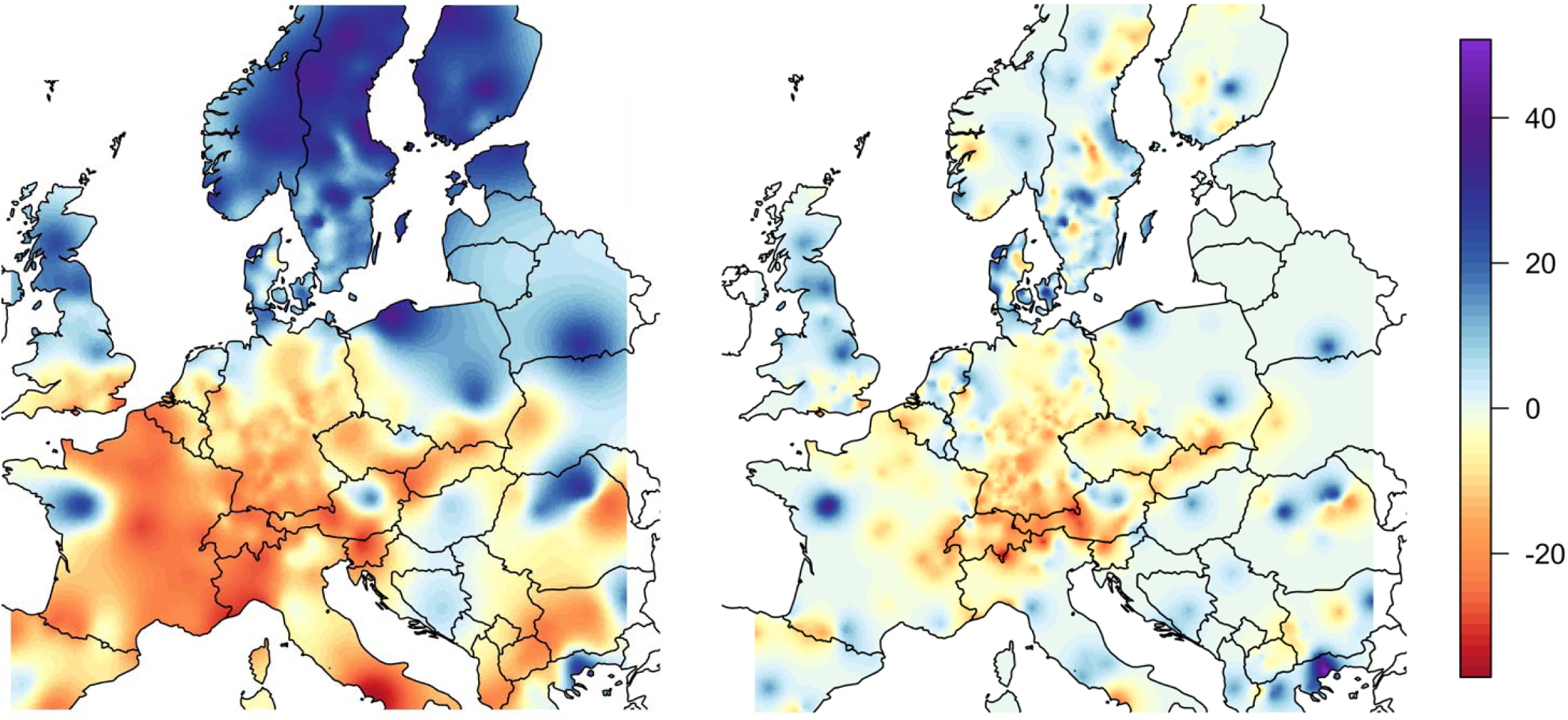
Spatial variation in flowering time [days] in model A (left) and model B (right) after the effects of the covariates (model A: year and elevation; model B: year, elevation, spring and winter temperature, and spring precipitation) have been accounted for

## Discussion

Herbaria are unique archives for studying long-term responses of plant phenology to anthropogenic climate change. Here, we studied herbarium specimens of 20 early-flowering forest herbs across Europe and show that these plants advanced their flowering during the last century, most likely in response to increasing spring temperatures. The herbarium data we used were substantially autocorrelated – even after accounting for elevation, climate and year, and including this spatial structure in our statistical models significantly improved the model fit and parameter estimates. Below, we therefore discuss only the results from models that accounted for spatial correlation.

### Temporal shifts in plant phenology

We found that forest understory herbs from Central Europe advanced their flowering by an average of six days during the last century (−0.6 days per decade). This is at the moderate end of what other studies found. Previous herbarium studies conducted in the temperate zone, which included 28-186 herbaceous or woody species and covered 100-170 years of data, estimated flowering time shifts between −0.4 and −1.5 days per decade (Primack et al. 2004, Miller-Rushing et al. 2006, Panchen et al. 2012, Molnar et al. 2012, Bertin 2015, Bertin et al. 2017). All of these studies were geographically very restricted and, except for one study from Hungary (Molnar et al. 2012), all came from the Northeastern US. There have been other longer-term studies on phenology trends in Europe, but these were based on field observations, and they did not go back further than the 1970s. The trends reported in these observational studies tend to be much stronger (−2.5 to −4.5 days per decade; Fitter and Fitter 2002, Menzel et al. 2006), possibly indicating that phenological changes have been accelerating during the last decades in response to more rapid climate changes (European Environmental Agency 2020). Interestingly, while herbarium studies from temperate regions were all relatively consistent, studies from other climatic regions found very different results, e.g. weaker or no phenology shifts across >1700 species in the subtropical southeastern US (Park and Schwartz 2015), or much stronger phenology shifts in some Himalayan species (up to −9 days per decade; Gaira et al. 2011, 2014). The stronger shifts in the Himalayas might at least be partly due to stronger climate changes at higher elevations, or due to greater temperature sensitivity of higher-elevation plants (see also discussion below).

### Relationships with climate warming

The long-term changes in plant phenology we detected are likely responses to climate change, in particular rising spring temperatures, which were strongly associated with the average collection dates of our herbarium specimens. For each 1°C of temperature increase, plants were on average collected −3.6 days earlier. In Europe, land temperatures have increased around 1.5°C since 1900 (Luterbacher et al. 2004, Harris et al. 2014, European Environmental Agency 2020), so the magnitude of overall phenological changes we observed is similar to what would be expected based on climate change and the observed temperature sensitivities (1.5°C x 3.6 days/°C = 5.4 days – vs. our observed average shift of around 6 days). Our results for temperature-phenology associations fit well to what others observed. Previous herbarium studies from the temperate zone estimated flowering-time advancements of −2.4 to −6.3 days per 1°C temperature increase (Primack et al. 2004, Miller-Rushing et al. 2006, Panchen et al. 2012, Calinger et al. 2013, Hart et al. 2014, Bertin 2015, Davis et al. 2015, Bertin et al. 2017). Again, most of these studies were from the Northeastern US, and they were often geographically very restricted. Two previous herbarium studies from Europe found stronger shifts of −6 to −13 days per 1°C (Robbirt et al. 2011, Diskin et al. 2012), but both were based on single species in rather restricted geographic areas. More robust European data comes from field observations: a long-term (1954-2000) observational study in England found advances of −1.7 to 6.0 days per 1°C across 385 plant species (Fitter and Fitter 2002), and a meta-analysis of long-term observation data found an average advancement of plant phenology of 2.5 days per 1°C temperature increase (Menzel et al. 2006). In a field monitoring study of a subset of 16 of this study’s species, we recently related plant phenology to forest microclimates and found a similar advancement of −4.5 days per 1°C temperature increase (Willems et al. 2021). So the overall pattern of 3-4 days earlier phenology per degree warming appears rather robust across a range of species and temperate regions, and our study strongly indicates that this large-scale biological response to anthropogenic climate change has also been taking place in Europe during the last century. As for the temporal shifts, our conclusions are restricted to temperate regions. Some studies from other climatic regions have found very different results, e.g. delayed rather than advanced flowering in response to increased spring temperatures in the subtropical southeastern US (Park and Schwartz 2015), or much stronger climate-related shifts in both directions in studies from Australia (Gallagher et al. 2009, Rawal et al. 2015). That the plants in our study also flowered earlier with warmer winter temperatures suggests that their potential chilling requirements (indicating that winter has passed) are yet still fulfilled – however if winter temperatures warm further if climate change intensifies this might change.

### Other drivers of phenology variation

While temperature may be a key driver of phenology, it is not the only one, and often does not explain all observed phenology variation (Marchin et al. 2015, Piao et al. 2019). In our study, we found that, across the study area, plants flowered later at higher elevation, and this pattern remained significant even if temperature was included as explanatory variable. Thus, the later flowering at higher elevations must be more than a temperature effect, and it indicates that phenology advances are generally slower at higher altitudes. One explanation could be that plants at higher elevation are less sensitive to temperature changes (Vitasse et al. 2010, Dai et al. 2014). On the other hand, the residual spatial variation we observed in model B indicates that in some mountainous regions (especially the Alps) plants flowered earlier than expected (after accounting for all covariates) and therefore, on the contrary, might be more sensitive to temperature changes (Chapman 2013, Liu et al. 2014). A solution for this apparent contradiction could be that the relationship between elevation and phenology is non-linear or is confounded with other environmental variables. Several other studies that related phenology to altitude provide mixed results, from slower to faster phenology changes at high elevations (Defila and Clot 2005, Ziello et al. 2009, Čufar et al. 2012, Kopp et al. 2020). Clearly, the relationship between elevation and phenology changes is not well understood yet, and large-scale herbarium plus climate data that correct for spatial autocorrelation have the potential to shed more light on this and to help to understand how, when, where and for which species elevation influences phenology.

Besides temperature, another climate factor that could potentially influence plant phenology is precipitation. We had expected a significant interaction with temperature, with strongest phenology advances where both temperature and precipitation were increasing, but there was no evidence for precipitation-phenology relationships in our data at all. Previous research found that changes in rainfall and water availability can influence phenology but with substantial geographical differences, e.g. in Mediterranean forests and shrublands (Peñuelas et al. 2004). Another complication with precipitation effects on phenology is that if precipitation occurs as snow this may influence phenology in very different ways than rain fall. Increased snow fall often delays plant growth and flowering (Park and Mazer 2018) – another potential explanation for why overall plants flowered later at higher elevations in our study. As global warming is expected to change snow melt more severely at higher elevations, it might have quite different effects on species at higher altitudes than on those at lower elevation (Cornelius et al. 2013), which in turn can cause problems for migrating or hibernating animal species across altitudinal gradients (Inouye et al. 2000).

### Spatial variation in phenology

Spatial autocorrelation has so far been largely ignored in herbarium-based studies of long-term phenology changes. However, it is important to take spatial variation into account not only because herbarium data are generally strongly spatially clustered, but also because neither phenology nor phenological responses to climate change are expected to be spatially homogenous across larger geographic scales. For previous studies that were geographically very restricted (Bertin 1982, Primack et al. 2004, Miller-Rushing et al. 2006, Miller-Rushing and Primack 2008, Bertin et al. 2017), the problem may be minor, but larger-scale analyses will require to take spatial variation into account. Recently, Park and Mazer (2018) studied phenological shifts across several climatic zones and Park et al. (2019) and Kopp et al. (2020) explicitly tested for geographic differences in phenological sensitivities in North America. To our knowledge, our study is the first herbarium-based study that modelled and mapped such spatial variation as a continuous variable in an analysis of large-scale phenology variation.

The most conspicuous pattern in the residual spatial variation of our data was that there appeared to be systematic differences in phenology associated with latitude, even in model B that accounted for climatic variation. In particular, plants from Central Europe (especially around the Alps) flowered earlier than predicted by our model. Such deviations indicate that we are either missing an important driver, or that plant responses to some of the covariates in the model differ geographically. Previous studies often found phenology shifts at high latitudes to be stronger in absolute terms (Root et al. 2005, Parmesan 2007, Ge et al. 2015) but weaker in relative terms (= per degree warming) than at low latitudes. This is usually explained by stronger temperature increases in northern regions (IPCC 2014), but lower temperature sensitivity of norther populations (Dai et al. 2014, Shen et al. 2015, Ge et al. 2015, Wang et al. 2015a, 2015b, Park et al. 2019, but see Wolkovich et al. 2012). The later may be a (late frost) risk-avoiding adaptation to variable, less reliable climates, causing plants populations to rely more on photoperiod (Renner and Zohner 2018). However, some studies also found that plants from (far) northern regions are more sensitive to temperature and require less warming to trigger leaf out or flowering (Riihimäki and Savolainen 2004, Pudas et al. 2008, Liang and Schwartz 2014, Prevéy et al. 2017), ensuring that plants start growing as soon as growth conditions become good in early spring, which may be crucial in cold regions with short growing seasons. Consequently, phenological sensitivity to temperature might decrease from southern to mid-northern latitudes but increase again in far-northern regions. This could indeed explain the earlier-than-expected phenology that we observed in Central Europe and the Alps. The missing consensus among studies about the association between latitude and phenology may be partially due to differences in spatial scale and because their relationship is complex, confounded with other environmental factors such as temperature, elevation or non-linear (Riihimäki and Savolainen 2004, Chmura et al. 2019, Kopp et al. 2020). These are challenges, that can be tackled by investigating geographic patterns via a continuous spatial field (as we did here, using R-INLA), that can depict differentiated geographic variability of phenology.

## Conclusions

The flowering time of forest herbs in Europe has substantially advanced during the last century, and these advances are strongly associated with climate warming. Our study demonstrates how herbarium specimens can be used to expand not only the temporal but also geographic and taxonomic scope of phenology research, and to contribute to understanding global environmental change (Wolkovich et al. 2014). Herbarium data from large geographic ranges are particularly powerful but they also come with challenges, and we showed that accounting for spatial autocorrelation significantly improved model fits and parameter estimates. Future studies should more frequently employ such spatial modeling techniques when analysing large-scale phenology variation and its different drivers, ideally across multiple climatic regions (Park et al. 2019).

The long-term phenology changes we observed in our study reflect physiological responses to climate warming, i.e. plants have adjusted to climate change (Munguía-Rosas et al. 2011). While this may to some extent be considered good news, the phenological shifts can have further consequences for the species and their associated ecological communities. For individual plant species, phenology shifts could be detrimental e.g. if they do not track warming temperatures well enough (Willis et al. 2008, Munguía-Rosas et al. 2011) or if earlier leaf-out or flowering increases the risk of late-frost damage (Wipf et al. 2006, Inouye 2008, Zohner et al. 2020). In addition, if climate change affects plants and their interacting organisms such as pollinators or herbivores unequally, then the phenology shifts of plants could result in temporal “mismatches” between interacting organisms (Renner and Zohner 2018). Finally, changes in plant phenology also influence ecosystem functions such as productivity or carbon cycling (Menzel et al. 2006, Cleland et al. 2007, Piao et al. 2019). Understanding not only phenology changes but also their further consequences for communities and ecosystems is an important goal for future research.

## Acknowledgements

We are very grateful for the help and support we received from many people working at the visited herbaria, especially Cornelia Dilger-Endrulat (Herbarium Tubingense), Frank Hellwig, Jochen Müller, Gabriele Reislöhner, Kirstin Victor (Herbarium Haussknecht, Jena) and Anette Rosenbauer, Mike Thiv, Arno Wörz (Herbarium Stuttgart). This work has been supported by the DFG Priority Program 1374 “Infrastructure-Biodiversity-Exploratories” (DFG project BO 3241/7-1 to OB). Further we thank the managers of the three Exploratories, Kirsten Reichel-Jung, Iris Steitz, and Sandra Weithmann, Juliane Vogt, Miriam Teuscher and all former managers for their work in maintaining the plot and project infrastructure; Christiane Fischer for giving support through the central office, Andreas Ostrowski for managing the central data base, and Markus Fischer, Eduard Linsenmair, Dominik Hessenmöller, Daniel Prati, Ingo Schöning, François Buscot, Ernst-Detlef Schulze, Wolfgang W. Weisser and the late Elisabeth Kalko for their role in setting up the Biodiversity Exploratories project. The authors have no conflict of interest to declare.

## Author contributions

FMW, OB and JFS designed the study; FMW collected the herbarium data; FMW compiled and analyzed all data; and FMW wrote the manuscript with all coauthors contributing to revisions.

