## Supplementary material for "Forest wildflowers bloom earlier as Europe warms – but not everywhere equally"

**Table S1:** The 20 studied forest understories herbs, their respective families and the number of herbarium specimens of each species that were included in the analyses.

| **Species** | **Family** | **N** |
| --- | --- | --- |
| *Adoxa moschatellina* | Adoxaceae | 415 |
| *Alliaria petiolata* | Brassicaceae | 252 |
| *Allium ursinum* | Amaryllidaceae | 206 |
| *Anemone nemorosa* | Ranunculaceae | 661 |
| *Anemone ranunculoides* | Ranunculaceae | 165 |
| *Arum maculatum* | Araceae | 46 |
| *Cardamine bulbifera* | Brassicaceae | 272 |
| *Euphorbia amygdaloides* | Euphorbiaceae | 93 |
| *Ficaria verna* | Ranunculaceae | 445 |
| *Galium odoratum* | Rubiaceae | 295 |
| *Lathyrus vernus* | Fabaceae | 343 |
| *Mercurialis perennis* | Euphorbiaceae | 349 |
| *Oxalis acetosella* | Oxalidaceae | 85 |
| *Paris quadrifolia* | Melanthiaceae | 409 |
| *Polygonatum verticillatum* | Asparagaceae | 64 |
| *Primula elatior* | Primulaceae | 193 |
| *Pulmonaria obscura* | Boraginaceae | 460 |
| *Ranunculus auricomus agg.* | Ranunculaceae | 330 |
| *Stellaria holostea* | Caryophyllaceae | 448 |
| *Viola reichenbachiana* | Violaceae | 600 |

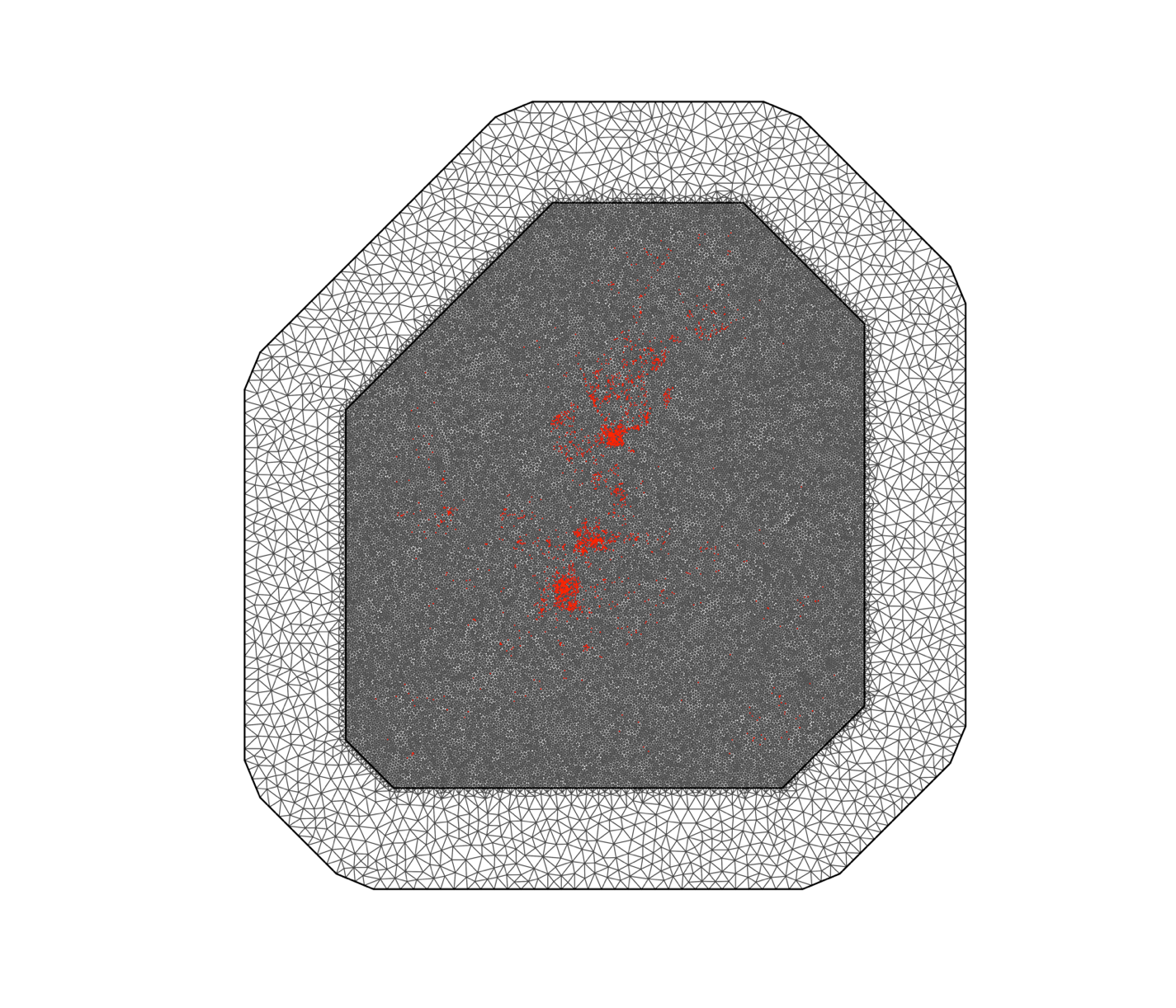

**Figure S1**: The mesh – based on refined Delaunay triangulation – used to estimate spatial autocorrelation of herbarium specimen data across Europe. The covariance matrix is estimated only for the inner area (see Methods section), and the outer area is a buffer zone against potential boundary effects. The sampling locations (red dots) are projected from latitude/longitude values onto the UTM coordinate system.

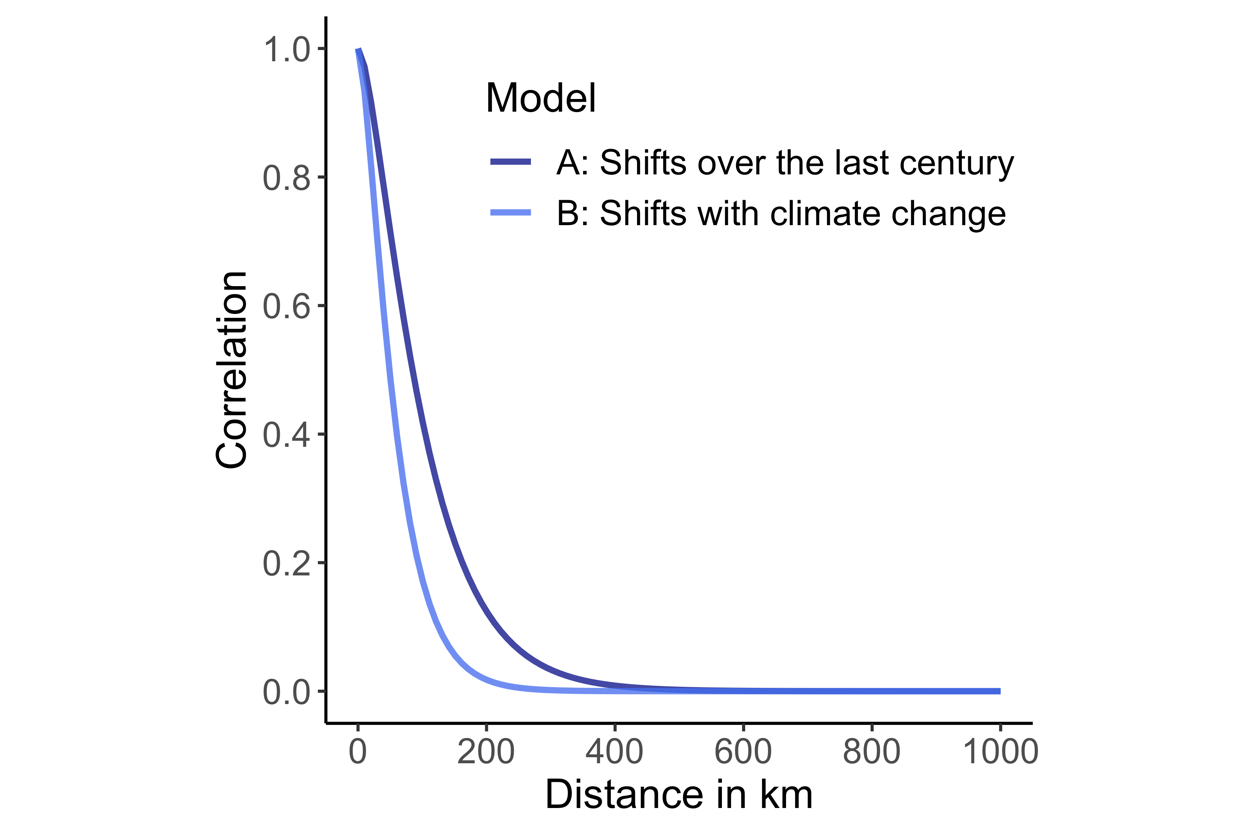

**Figure S2.** The spatial correlation of flowering time data in models A and model B (after the effect of the covariates have been accounted for).

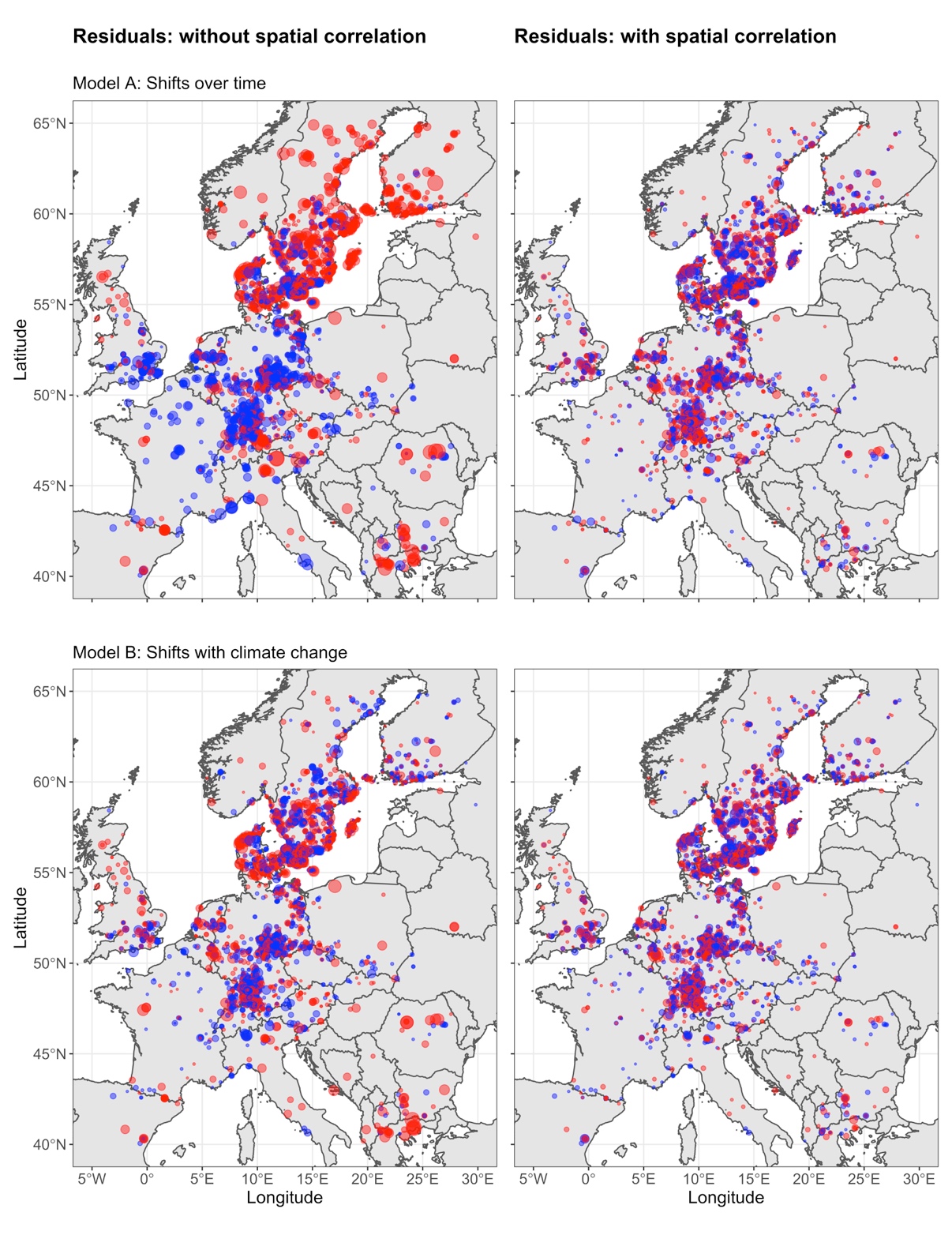

**Figure S3.** Spatial patterns in the residuals of the model A (top panels) and model B (bottom panels), without (left) and with (right) accounting for spatial correlation in the models. Larger points indicate larger residuals and point color indicates whether residuals were negative (red) or positive (blue).

**Model Validation Check for Model A: Phenological shifts over the last century**

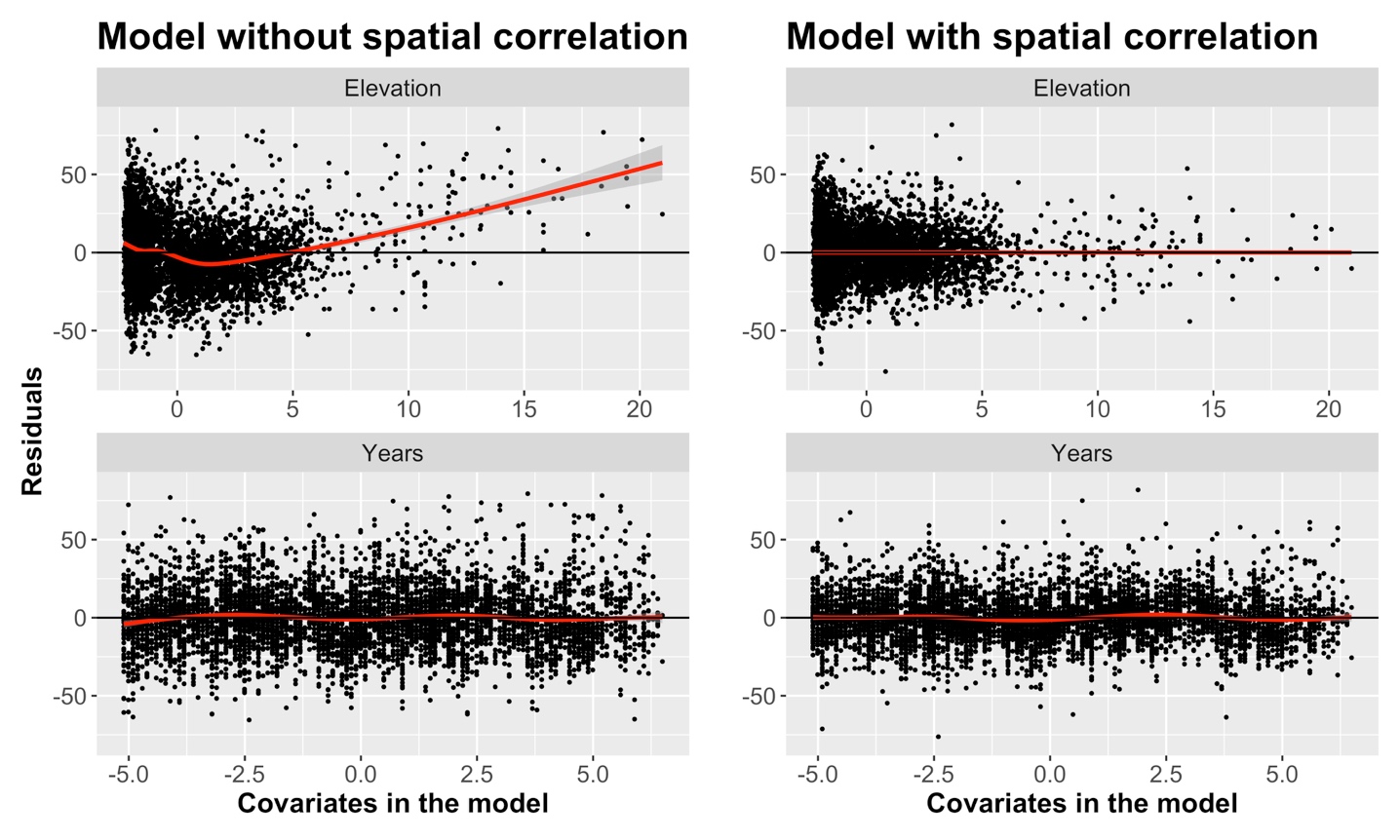

**Figure S4.** Residuals plotted against each covariate in model A without (left) and with spatial correlation (right). The red lines are thin plate regression spline smoothers, fitted with the *mgcv* package (Wood 2006), with 95% confidence intervals in gray, to aid visual interpretation.

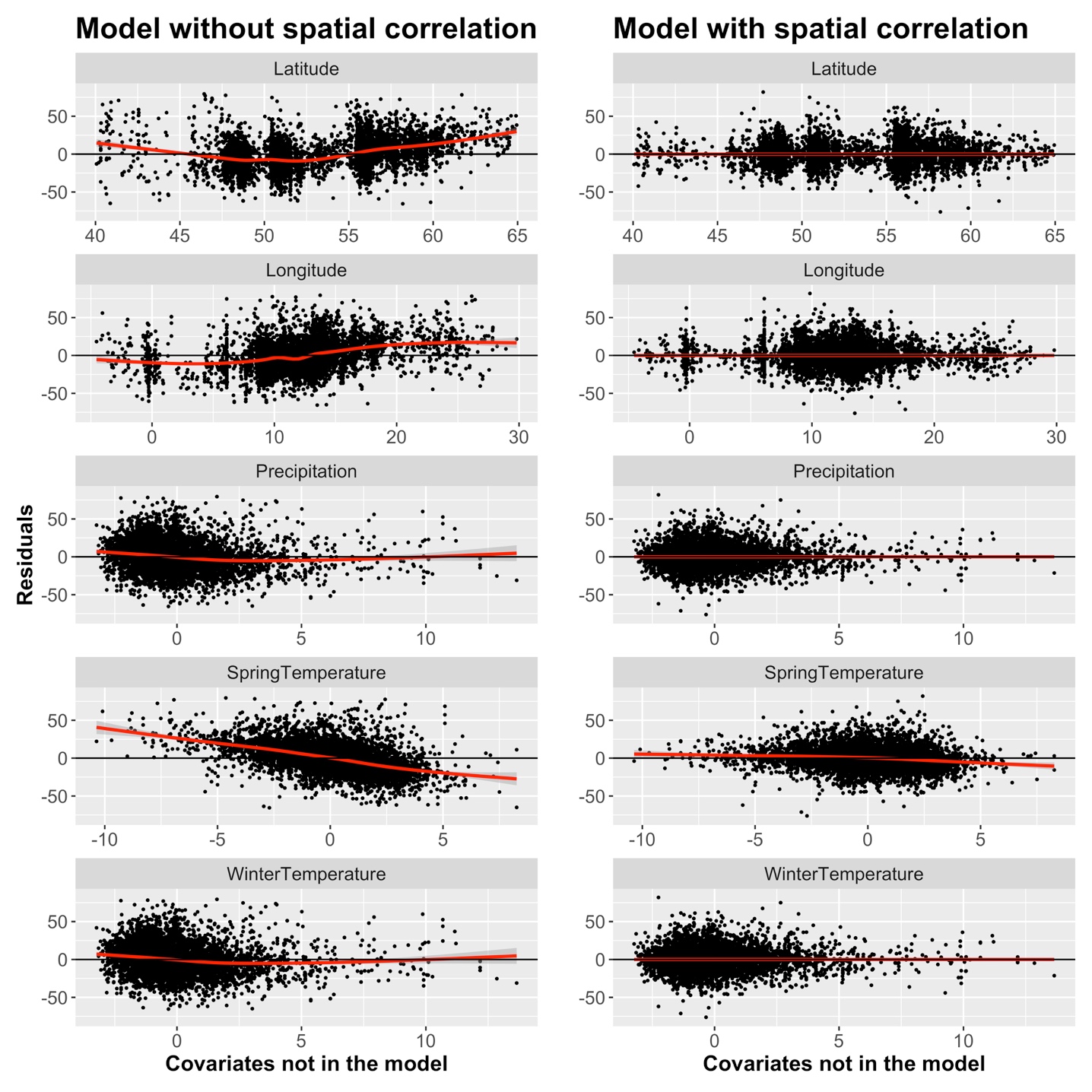

**Figure S5.** Residuals of Model A plotted against covariate that were not included in model A, without (left) and with spatial correlation (right). The red lines are thin plate regression spline smoothers, fitted with the *mgcv* package (Wood 2006), with 95% confidence intervals in gray, to aid visual interpretation.

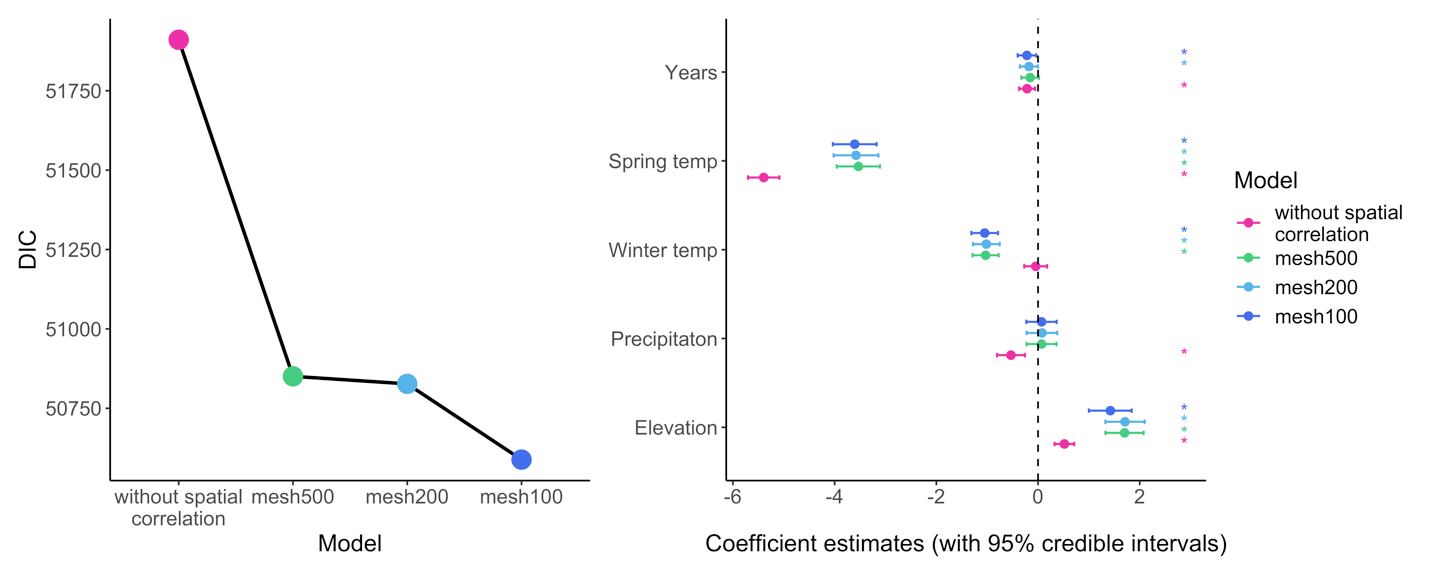

**Figure S6.** DIC values and model estimates (regression coefficients) of model B without spatial correlation and with spatial correlation, using different mesh sizes. For our analyses with spatial correlation we chose the finest mesh100 (number indicates the assumed range of spatial correlation).

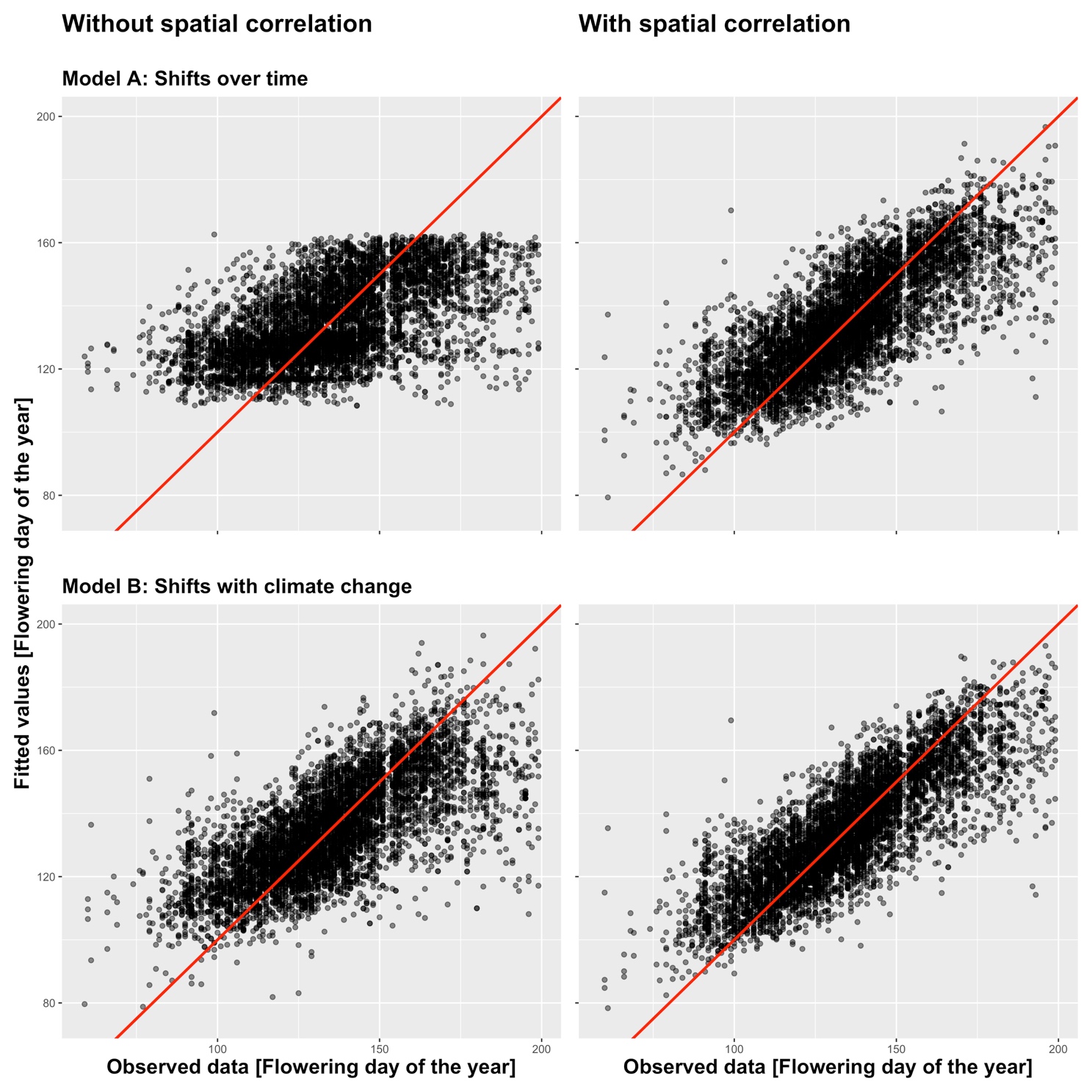

**Figure S7.** Observed vs. fitted values for model A (top) and model B (bottom), without (left) and with (right) spatial correlation. The red diagonal is the identity line. In both models that included spatial correlation the fitted values were much closer to the observed values (r = 0.78 vs. 0.57 for Model A and r = 0.82 vs 0.70 for Model B; Fig. S7).

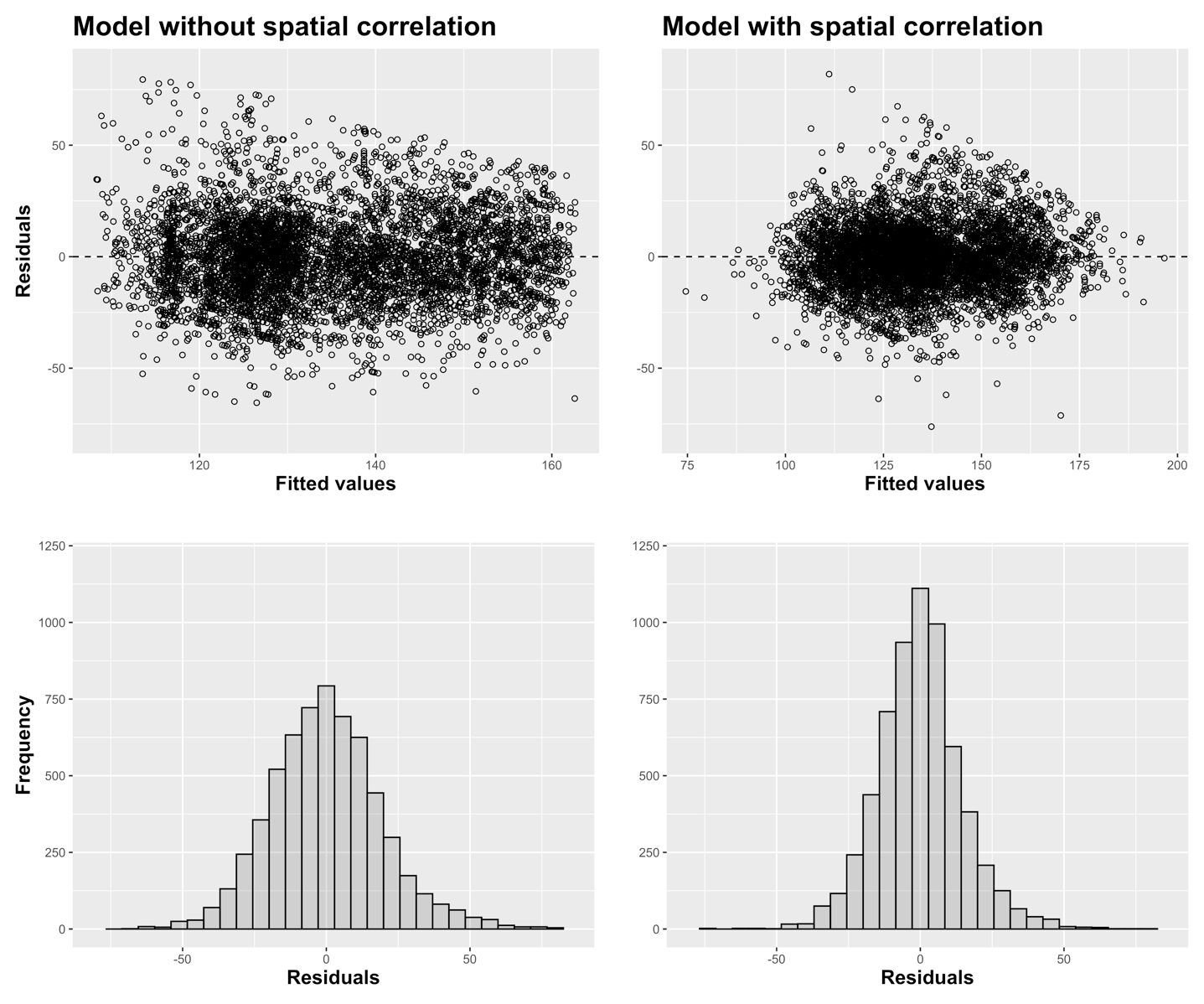

**Figure S8**: Residuals vs. fitted values (top) and histogram of the residuals (bottom) of model A without (left) and with spatial correlation (right). Residuals and fitted values of model B showed similar patterns.

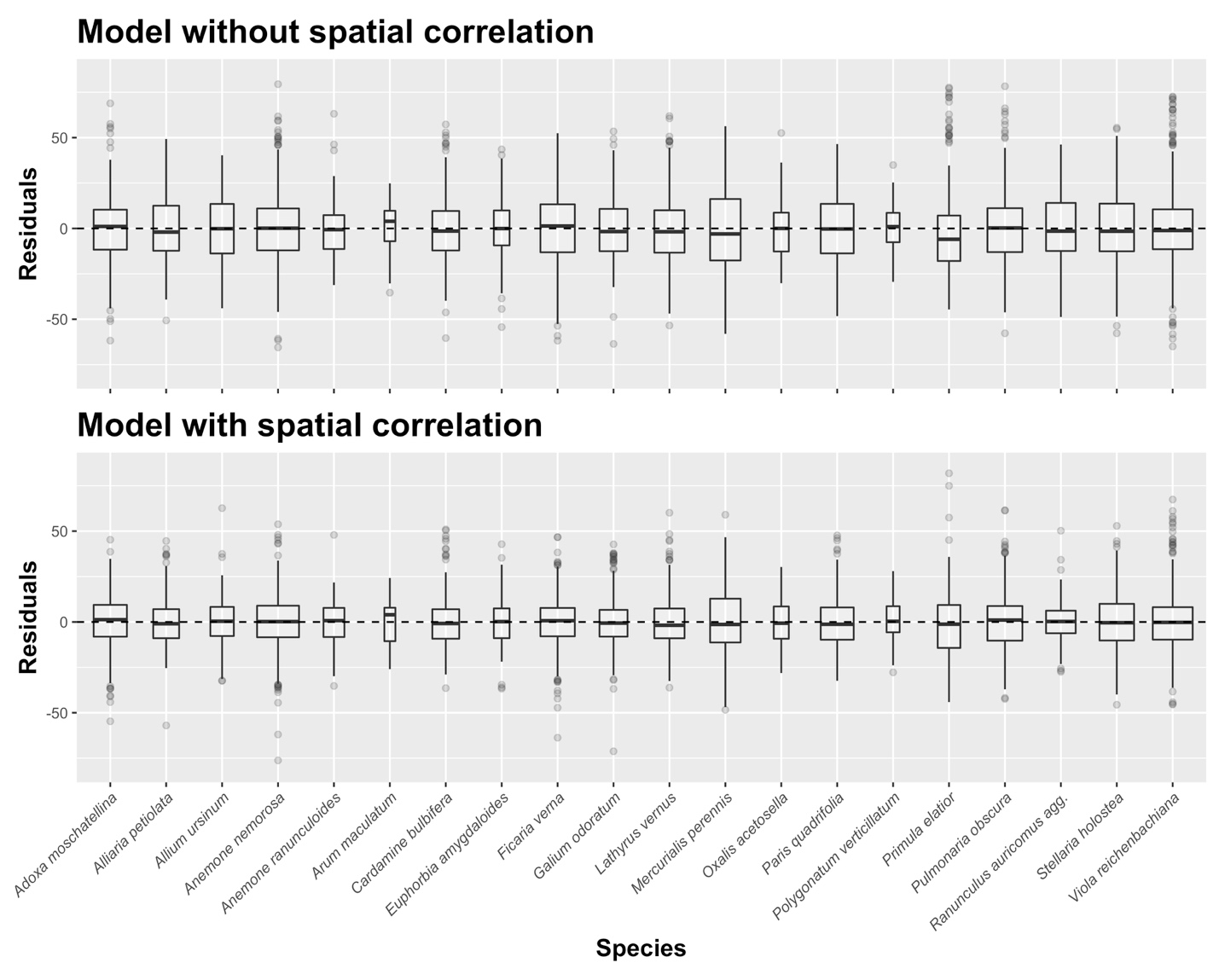

**Figure S9**: Residuals (of model A) for each species (that were included as a random factor) without (top) and with spatial correlation (bottom). Box widths are proportional to sample size.

**Table S2.** Estimates, standard deviations and 95% credible intervals for the hyperparameter values in model A with spatial correlation. RE = random effect.

|  | **Estimate** | **SD** | **95% CI** |
| --- | --- | --- | --- |
| Precision for the Gaussian observations | 4.27E-03 | 1.00E-04 | 4.09E-03, 4.48E-03 |
| Precision for RE species intercepts | 8.47E-03 | 3.99E-03 | 3.72E-03, 1.88E-02 |
| Precision for RE species slopes | 1.83E+04 | 1.93E+04 | 2.07E+03, 6.91E+04 |
| Range for spatial RE [km] | 214.07 | 64.94 | 92.06, 331.46 |
| SD for spatial RE [m] | 1.88 | 1.89 | 14.8, 22.11 |

**Table S3.** Model estimates, standard deviations and 95% credible intervals for the parameters in model A without spatial correlation. RE = random effect.

|  | **Estimate** | **SD** | **95% CI** |
| --- | --- | --- | --- |
| Intercept | 135.10 | 2.55 | 130.09, 140.10 |
| Years [Decades] | -1.43 | 0.09 | -1.60, -1.25 |
| Elevation [100 m] | -0.34 | 0.10 | -0.53, -0.14 |
| Precision for the Gaussian observations | 2.56E-03 | 4.07E-05 | 2.49E-03, 2.65E-03 |
| Precision for RE species intercepts | 7.86E-03 | 2.98E-04 | 7.29E-03, 8.48E-03 |
| Precision for RE species slopes | 1.58E+04 | 5.41E+02 | 1.48E+04, 1.70E+04 |

**Table S4**. Estimates, standard deviation and 95% credible intervals for the parameters and hyperparameters in the model B version without spatial correlation. RE = random effect.

|  | **Estimate** | **SD** | **95% CI** |
| --- | --- | --- | --- |
| Intercept | 135.90 | 2.50 | 130.94, 140.77 |
| Spring Temperature [°C] | -0.21 | 0.08 | -0.37, -0.06 |
| Winter Temperature [°C] | -5.39 | 0.16 | -5.70, -5.09 |
| Precipitation [mm/10] | -0.53 | 0.14 | -0.81, -0.26 |
| Elevation [100 m] | 0.52 | 0.10 | 0.33, 0.71 |
| Years [Decade] | -0.05 | 0.11 | -0.27, 0.18 |
| SpTemp:Year | 0.06 | 0.04 | -0.01, 0.13 |
| SpTemp:Elevation | 0.13 | 0.04 | 0.06, 0.20 |
| SpTemp:Precipitation | -0.04 | 0.05 | -0.15, 0.06 |
| SpTemp:WinterTemperature | -0.07 | 0.02 | -0.12, -0.02 |
| Precision for the Gaussian observations | 4.0e-03 | <0.001 | 4.00e-03, 4.00e-03 |
| Precision for RE species intercepts | 7.0e-03 | 2.00e-03 | 4.00e-03, 1.10e-03 |
| Precision for RE species slopes | 4.8e+04 | 1.52e+04 | 2.56e+04, 8.45e+04 |

**Table S5**. Estimates, standard deviations and 95% credible intervals for the hyperparameter values in model B with spatial correlation. RE = random effect.

|  | **Estimate** | **SD** | **95% CI** |
| --- | --- | --- | --- |
| Precision for the Gaussian observations | 0.0047 | 0.0001 | 0.0046, 0.0048 |
| Precision for RE species intercepts | 0.0089 | 0.0033 | 0.0043, 0.0170 |
| Precision for RE species slopes | 481.9 | 361.4 | 76.5, 1414.8 |
| Range for spatial RE [km] | 113.61 | 20.64 | 75.13, 162.53 |
| SD for spatial RE [m] | 16.23 | 1.23 | 13.70, 18.94 |

1.2336
